# Systematic Identification of Correlates of HIV-1 Infection: An X-Wide Association Study in Zambia

**DOI:** 10.1101/126052

**Authors:** Chirag J. Patel, Jay Bhattacharya, John P.A. Ioannidis, Eran Bendavid

## Abstract

**Background:** HIV-1 remains the leading cause of death among adults in Sub-Saharan Africa, and over 1 million people are infected annually. Better identification of at-risk groups could benefit prevention and treatment programmes. We systematically identified factors related to HIV-1 infection in two nationally representative cohorts of women that participated in Zambia’s Demographic and Health Surveys (DHS).

**Methods:** We conducted a comprehensive analysis to identify and replicate the association of 1,415 social, economic, environmental, and behavioral indicators with HIV-1 status. We used the 2007 and 2013-2014 DHS surveys conducted among 5,715 and 15,433 Zambian women, respectively (727 indicators in 2007; 688 in 2013-2014; 688 in both). We used false discovery rate criteria to identify indicators that are strongly associated with HIV-1 in univariate and multivariate models in the entire population, as well as in subgroups stratified by wealth, residence, age, and history of HIV-1 testing.

**Findings:** In the univariate analysis we identified 102 and 182 variables that are associated with HIV-1 in the 2007 and 2013-2014 surveys, respectively, among which 79 were associated in both. Variables that were associated with HIV-1 status in all full-sample models (unadjusted and adjusted) as well as in at least 17 out of 18 subgroups include being formerly in a union (adjusted OR 2007 2.8, p<10^−16^; 2013-2014 2.8, p<10^−29^), widowhood (adjusted OR 2007 3.7, p<10^−12^; 2013-2014 4.2, p<10^−30^), history of genital ulcers in the last 12 months (adjusted 2007 OR 2.4, p<10^−5^; 2013-2014 2.2, p<10^−6^), and having a woman for the head of the household (2007 OR 1.7, p<10^−7^; 2013-2014 OR 2.1, p<10^−26^), while owning a bicycle (adjusted 2007 OR 0.6, p<10^−6^; 2013-2014 0.6, p<10^−8^) and currently breastfeeding (adjusted 2007 OR 0.5, p<10^−9^; 2013-2014 0.4, p<10^−26^) were associated with decreased risk. Using the identified variables, area under the curve for HIV-1 positivity ranged from 0.76 to 0.82.

**Interpretation:** Our X-wide association study in Zambian women identifies multiple under-recognized factors correlated with HIV-1 infection in 2007 and 2013-2014, including widowhood, breastfeeding, and being the head of the household. These variables could be used to improve HIV-1 testing and identification programs.

## Introduction

Approaching the public health goals for global HIV-1 such as “90-90-90” (90% of those with HIV-1 aware of their status, 90% in regular treatment, and 90% of those on treatment virally suppressed) requires effective identification of at-risk and infected individuals.^1,2^ Despite large efforts to expand testing, treatment, and retention services, only 45% of HIV-infected individuals in Sub-Saharan Africa knew their HIV-1 status in 2013, and estimated antiretroviral therapy (ART) coverage in 2014 exceeded 50% of all those infected in only five countries.^3-6^ A potential approach to improving HIV-1 testing and diagnosis is to better target individuals and populations for testing and care. Existing HIV-1 control programs, such as the US President’s Emergency Plan for AIDS Relief, increasing use data-driven approaches to align resources towards high-burden populations.^7,8^

Current understanding of HIV-1 risk factors in Sub-Saharan Africa commonly come from nationally representative surveys such as the Demographic and Health Surveys (DHS) that include HIV-1 testing and report prevalence stratified by pre-specified groups such as age, education, place of residence, and number of sexual partners.^9-11^ Such HIV-1 testing and epidemiologic stratification was carried out in two DHS surveys conducted in Zambia in 2007 and again in 2013-2014 in nationally representative samples of 5,715 and 15,433 women, respectively.^12,13^ However, selective identification of risk factors by testing one or only a few factors at a time may lead to incomplete understanding of or even misleading notions about possible risk factors.^14,15^ Traditional risk factors (age, gender, education, place of residence, and number of sexual partners) explain less than 10% of the variation in HIV-1 infection, and represent only a small fraction of the information available in the surveys.^12,13^ While risk factors such as age and gender are intuitive and important, unintuitive or under-recognized correlates that may identify novel high-risk groups and generate new hypotheses for further study and intervention design.

We present an approach for systematically assessing the relationship between HIV-1 status and many putative risk factors. We exploit the breadth of DHS surveys to conduct an *X*-wide association study (XWAS) of HIV-1 risk, where *X* stands for all social, behavioral, environmental, and economic factors that are reliably available in DHS. This approach systematically associates each available variable with HIV-1 status, as is done in current-day genomics investigations (e.g. genome-wide association studies [GWASs]). We have previously utilized the approach to systematically study the association of environmental exposures, dietary indicators, clinical biomarkers, and micronutrient blood tests associated with outcomes such as type II diabetes, blood pressure, mortality, and income.^16-19^ An advantage of this approach is that variables are examined using a systematic approach, thus avoiding selective reporting bias, while controlling for the rate of false positives.^20,21^

## Methods

### Overview

We used the 2007 and 2013-2014 DHS surveys from Zambia, where HIV-1 prevalence among women 15 to 40 years old was estimated at 16.1% and 15.1%, respectively.^13^ We linked the HIV-1 test results with all the indicators in the individual women’s surveys. We split the data in each survey into training (“discovery”) and replication data, analyzed the association of each variable with HIV-1 status in univariate and adjusted analyses, and examined the stability of the findings over time and in population subgroups. Figure 1 illustrates the analysis steps using the 2013-2014 survey metadata.

**Figure 1:**
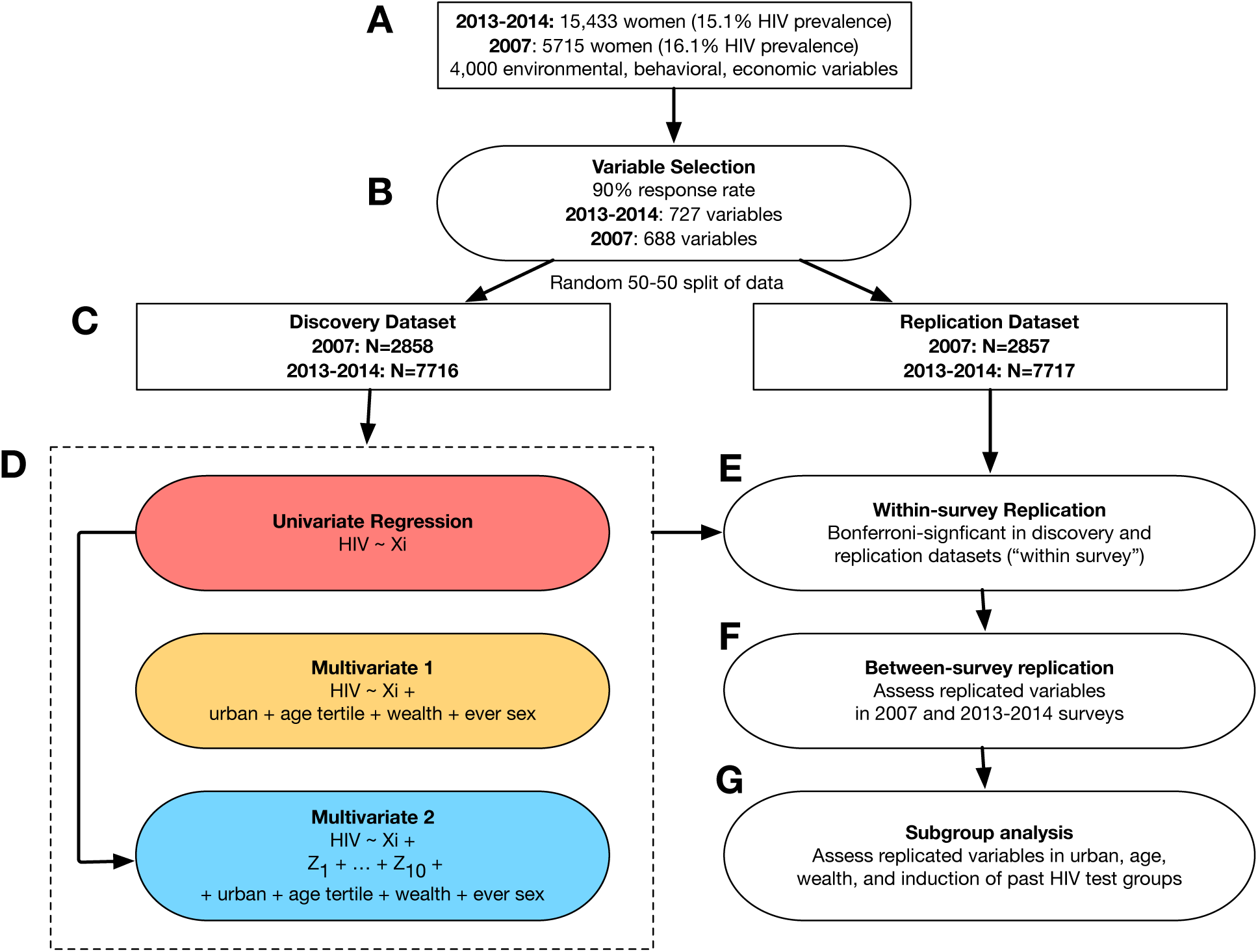
Schematic overview of XWAS process. A.) The Demographic and Health Study consists of 15,433 women (15% HIV-1 prevalence) in 2013-2014 and 5,715 women (16.1 prevalence) in 2007. B.) We selected variables that had >= 90% response rate and were non-redundant resulting in 688 total variables in both the 2007 and 2013-2014 surveys. C.) Split the data into two random subsets for discovery and replication (N=7,716 and 7,717 respectively for 2013-2014 and 2,858 and 2,857 for 2007). D.) We ran 3 model configurations: a univariate (red); a multivariate with adjustment variables selected a priori (yellow), including age, urban resident, wealth index, and ever had sex prior to survey; and a multivariate model consisting of variables identified in the univariate analysis (blue). X*i* denotes the *i*th variable out of 688 in 2013-2014 and 727 variables in 2007 respectively (688 overlapping). E.) We attempted to replicate results within surveys from models in the independent replication dataset. F.) We identified variables replicated between the 2007 and 2013-2014 surveys. G.) We executed subgroup analysis for each variable identified in the univariate regression.

### HIV-1 Status

The HIV-1 testing procedure in the surveys involved identifying eligible household members, obtaining consent, collecting dried blood spots, and processing and testing in a centralized lab. Two separate enzyme-linked immunosorbent assay (ELISA) tests were used for screening and confirmation of HIV-positive tests, with additional Western Blot confirmation for discordant ELISA tests. In both surveys, every test was definitively identified as positive or negative (i.e. there were no indeterminate tests). The HIV-1 test results were then linked one-to-one with the individual survey data.

### Selection of Social, Behavioral, Environmental, and Economic Indicators

We used the following process to identify and create the variables for the XWAS (Table 1 includes the variable selection process’ metadata). Starting with the raw data after removal of placeholder variables (e.g. birthdates of children 6-20 for mothers with 5 children), we recast all variables with 30 or fewer levels as binary variables. Variables with 30 or more levels were treated as continuous. This decision rule preserved most meaningfully continuous variables while discretizing most non-ordinal variables. Sensitivity analyses with thresholds ranging from 25 to 35 levels did not alter our findings. Then, we kept only those variables with at least 90% complete data to avoid what some consider unacceptable levels of missingness.^22^ This led to dropping over 40% of the variables in each survey. We removed variables with no variation (e.g. an indicator variable for completion of the survey), and kept the first occurrence in any pair of collinear variables with correlation coefficient>=0.99. The entire set of variables is in Supplementary Table 1.

**Table 1:** Variable Preparation Process

|  | **<u>2007 Survey</u>** |  | **<u>2013-2014 Survey</u>** |  |
| --- | --- | --- | --- | --- |
| *Variable List Preparation Stage* | *Change (n)* | *Remaining (n)* | *Change* | *Remaining* |
| Initial dataset |  | 976 |  | 957 |
| Discretize categorical variables | 705 | 1,681 | 681 | 1,638 |
| Remove variables with more than 10% missingness | -676 | 1,005 | -667 | 971 |
| Remove indicator and no-variation variables | -84 | 921 | -82 | 889 |
| Remove one from any pair of variables that are 99% or more correlated | -194 | 727*<br>(Final sample) | -201 | 688*<br>(Final sample) |

\* In the 2007 survey, 160 of the 727 variables in the final sample (22%) were unchanged from the original dataset. In the 2003-2014 survey, 160 (23%) were unchanged from the original dataset.

### Association Study Procedures

We divided each survey (5,715 women in 2007 and 15,433 women in 2013-2014), randomly into two equally sized (±1) datasets for discovery and replication. We conducted three XWAS analyses in the discovery dataset (Figure 1D): (i) a univariate analysis; (ii) an analysis adjusted for known HIV-1 risk factors (*ex ante* analysis); and (iii) an analysis that, in addition to the *ex ante* factors, adjusted for the ten variables that explained the greatest portion of the variation in HIV-1 status in the univariate analysis (*ex post* analysis).

In the first step we estimated univariate logistic regression models of the following form:

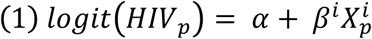

Where *HIV_p_* represents the HIV-1 status of person *p*, and 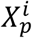 denotes the *i*th variable for person *p*. This procedure is repeated for each of the variables in the 2007 and 2013-2014 surveys. The exponentiation of *β^i^* corresponds to the odds ratio for HIV-1 per unit change for each variable *i*. To control for multiple hypothesis testing we calculated the Benjamini-Hochberg (BH) false discovery rate (FDR), the estimated proportion of discoveries made that were false.^23^ The BH method assumes independence between statistical tests and therefore counts correlated variables as independent for determining the discovery threshold (median absolute correlation between all pairs of replicated variables 0.06 (IQR 0.02 to 0.12) in 2007 and 0.04 (IQR, 0.02 to 0.09) in 2013). In all analyses, we used the survey HIV-1 sampling weights and Huber-White robust standard errors.^24^

In the *ex ante* analysis, we adjusted for five pre-determined (*ex ante*) controls: urban or rural residence, DHS wealth index (a discretized 5-level scale from poorest to wealthiest quintile, with poorest as reference), age, whether or not the respondent indicated she had previously been tested for HIV, and whether or she indicated that she never had sex at the time of the interview.^25^ Specifically, the model was implemented as follows:

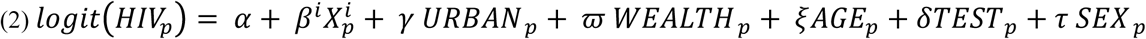
Where covariates for urban residence, wealth, age, past testing, and sexual debut are indexed by person (*p*), and 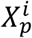 again denotes the *i*th exposure variable for person *p*.

In the *ex post* analysis, we adjusted for the ten variables that had the highest explanatory power (using Nagelkerke R^2^) among those replicated in the univariate analysis. The purpose of this analysis was to improve the identification of strong correlates that identify HIV-1 status even after controlling for the most explanatory variables.^26^ The ten variables were selected separately in the 2007 and 2013-14 surveys.

We had two levels of replication, within-survey replication and between-survey replication (Figure 1EF). We deemed a within-survey “replicated finding” for *β^i^* as one that had FDR<5% in the discovery dataset and the sign for *β^i^* was in the same direction in the replication dataset with a nominal p-value<0.05 (Figure 1E). The second level of replication is between-survey replication where we sought within-survey replication in both the 2007 and 2013-2014 surveys (Figure 1F).

Next, we assessed the predictive capability of HIV positivity of the variables found in all three modeling scenarios. For example, if variables *X^a^, X^b^, X^c^* were tentatively replicated in the univariate modeling scenario in the 2007 survey, we fit a model:

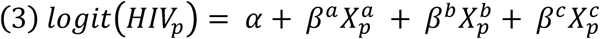
and assessed the Nagelkerke R^2^ and the area under the curve for the model.

We then assessed pairwise correlations among all of the replicated variables to assess the clustering of HIV-1 risk factors and variables. That is, we wanted to identify the clusters of variables that potentially measure a latent HIV-1 risk factor (e.g. if a host of household possession variables are all related to HIV-1 and are correlated among themselves, that may indicate that wealth, a likely latent variable they measure, is a risk factor for HIV). We visualized pairwise correlations in a heatmap.^27,28^

Finally, we tested the association of all replicated univariate findings in 9 subgroups (Figure 1G): (1-3) three age bins (15-<23; 23-<33; and 33-49); (4-5) two wealth groups (wealth quintiles 1-3 [poorer] and wealth quintiles 4-5 [wealthier]); (6-7) two residence groups (urban and rural); and (8-9) two groups based on whether or not the respondent indicated that they had ever received an HIV-1 test.

To promote reproducibility of this work, the analytic code is available in a Github repository, and the figures can be accessed at www.chiragjpgroup.org/hiv_zambia; all analyses were performed using Stata 14 (Statacorp) and R v3.2.2 (http://cran.r-project.org/). The sponsor of the study had no role in study design, data collection, data analysis, data interpretation, or writing of the report.

## Results

Our surveys included information on 5,715 Zambian women with HIV-1 test results in 2007 and 15,433 in 2013-2014. In the univariate analysis, 102 (out of 727, 14%) variables were replicated and associated with HIV-1 in 2007, and 182 (out of 688, 26%) in 2013-2014. Figure 2 shows a plot of p-values versus odds ratios of the association with HIV-1 of the variables tested in 2007 and 2013-2014. A total of 79 variables were associated with HIV-1 status in the univariate analysis, 30 variables in the *ex ante* analysis, and 8 variables in the *ex post* analysis in *both* 2007 and 2013-2014. Table 2 shows all the variables that were replicated *in both surveys* in at least one analysis. All replicated variables (in any analysis) appear in Supplementary Tables SA1.2-SA1.4. Supplementary Table SA1.5 shows the associations between the control variables and HIV in the *ex post* models.

**Figure 2:**
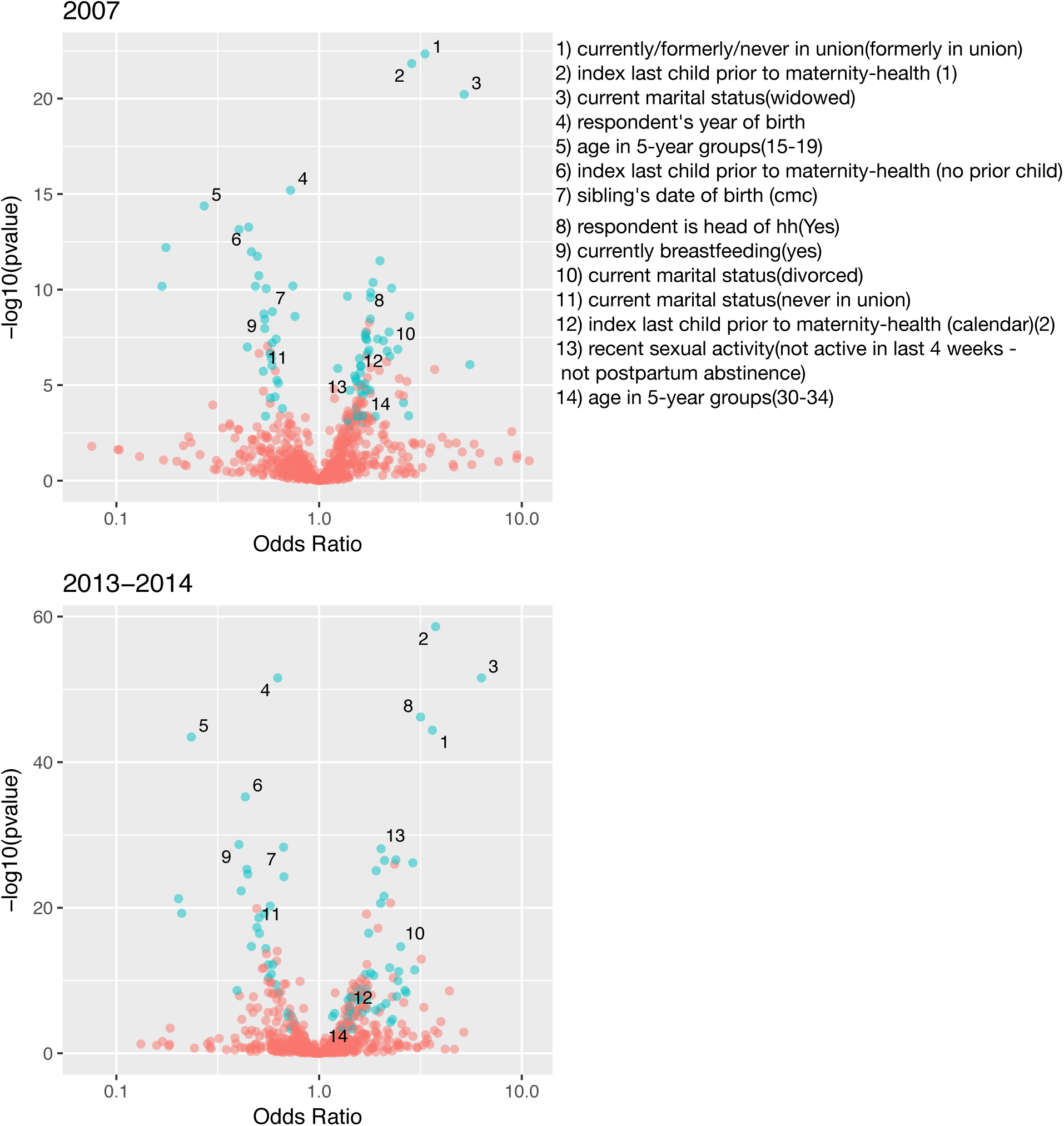
Volcano plot from univariate analysis depicting odds ratio versus −log10(pvalue) of association. Teal points denote replicated findings in each dataset (‘07 and ‘13-14).

**Table 2:** Univariate and adjusted associations with HIV-1 status

| Variable name | Value | *Univariate* |  | *Ex ante* |  | *Ex post* |  |
| --- | --- | --- | --- | --- | --- | --- | --- |
|  |  | OR<br>( '07; '13-4) | -log10(p)<br>( '07; '13-4) | OR<br>( '07; '13-4) | -log10(p)<br>( '07; '13-4) | OR<br>( '07; '13-4) | -log10(p)<br>( '07; '13-4) |
| desire for more children | wants after 2+ years | 0.5; 0.4 | 10.7*; 24.6* | 0.5; 0.5 | 7.9*; 13.1* | 0.5; 0.5 | 6.9*; 11.7* |
| currently breastfeeding | yes | 0.5; 0.4 | 8.7*; 28.7* | 0.5; 0.4 | 9.5*; 26.4* | 0.5; 0.3 | 6.9*; 22.3* |
| number of children 5 and under in household (de jure) | 1 | 1.7; 2.0 | 7.4*; 20.6* | 1.7; 2.0 | 6.8*; 15.4* | 1.9; 2.3 | 5.6*; 17.3* |
| currently/formerly/never in union | formerly in union/living with a man | 3.3; 3.6 | 22.3*; 44.4* | 2.8; 2.8 | 15.7*; 29.0* | -; 2.7 | -; 18.5* |
| index last child prior to maternity-health (calendar) | 1 | 2.9; 3.8 | 21.8*; 58.6* | 2.2; 2.6 | 9.1*; 22.6* | 1.8; 2.6 | 3.6; 15.0* |
| current marital status | widowed | 5.2; 6.3 | 20.2*; 51.6* | 3.7; 4.2 | 12.6*; 31.0* | 1.7; 4.5 | 1.3; 22.6* |
| respondent's line number | 1 | 2.3; 3.2 | 10.1*; 46.2* | 1.9; 2.4 | 5.7*; 27.3* | 0.9; 2.3 | 0.1; 19.5* |
| number of living children | 1 | 1.8; 1.3 | 9.6*; 2.9* | 1.7; 1.6 | 6.3*; 4.9* | 1.4; 1.6 | 2.4; 4.4* |
| number of children 5 and under in household (de jure) | 3 | 0.5; 0.6 | 8.4*; 10.9* | 0.6; 0.6 | 7.0*; 8.7* | 0.6; 0.6 | 3.3; 6.5* |
| current contraceptive method | condom | 2.2; 3.0 | 7.8*; 11.5* | 1.8; 2.5 | 4.6*; 8.5* | 1.5; 2.4 | 1.4; 5.9* |
| household has: bicycle | yes | 0.6; 0.6 | 7.4*; 9.5* | 0.6; 0.6 | 6.2*; 8.1* | -; 0.7 | -; 4.6* |
| sex of household head | female | 1.7; 2.1 | 7.4*; 26.5* | 1.7; 2.1 | 7.1*; 26.1* | 0.9; 2.1 | 0.2; 20.0* |
| current marital status | divorced | 2.2; 2.5 | 6.8*; 14.7* | 1.9; 1.9 | 4.6*; 8.0* | 0.8; 1.9 | 0.4; 5.6* |
| number of household members (listed) | 2 | 2.2; 2.5 | 6.5*; 10.0* | 2.2; 2.3 | 6.6*; 7.3* | 2.3; 2.6 | 4.3; 7.4* |
| age in 5-year groups | 15-19 | 0.3; 0.2 | 14.4*; 43.5* | 0.5; 0.5 | 4.3; 6.9* | 0.5; 0.5 | 2.6; 5.2* |
| daughters who have died | 1 | 2.0; 1.7 | 11.5*; 7.4* | 1.8; 1.2 | 7.4*; 1.3 | 2.3; 1.2 | 8.3*; 0.5 |
| unmet need | never had sex | 0.2; 0.2 | 10.2*; 19.2* | 0.4; 0.8 | 0.7; 0.1 | -; - | 42.5*; 89.5* |
| had genital sore/ulcer in last 12 months | yes | 2.8; 2.4 | 8.6*; 7.8* | 2.4; 2.2 | 5.9*; 6.1* | 1.8; 2.0 | 1.8; 2.9 |
| sons who have died | 0 | 0.6; 0.6 | 7.2<sup>*</sup>; 12.2<sup>*</sup> | 0.7; 0.8 | 3.7<sup>*</sup>; 1.8 | 0.6; 0.8 | 5.0<sup>*</sup>; 0.9 |
| recent sexual activity | not active in last 4 weeks - not postpartum abstinence | 1.5; 2.0 | 5.3<sup>*</sup>; 28.1<sup>*</sup> | 1.2; 1.8 | 0.9; 17.4<sup>*</sup> | 0.9; 1.8 | 0.4; 13.3<sup>*</sup> |
| desire for more children | wants within 2 years | 1.6; 2.4 | 4.5<sup>*</sup>; 26.6<sup>*</sup> | 1.6; 2.2 | 3.9; 19.8<sup>*</sup> | 1.7; 2.3 | 3.3; 15.0<sup>*</sup> |
| number of sex partners, excluding spouse, in last 12 months | 0 | 0.6; 0.6 | 4.4<sup>*</sup>; 12.2<sup>*</sup> | 0.6; 0.5 | 2.8; 12.9<sup>*</sup> | 0.9; 0.5 | 0.2; 9.7<sup>*</sup> |
| number of living children | 0 | 0.5; 0.5 | 11.7<sup>*</sup>; 18.6<sup>*</sup> | 0.9; 1.3 | 0.2; 2.0 | 1.3; 1.5 | 1.0; 3.3<sup>*</sup> |
| daughters who have died | 0 | 0.5; 0.6 | 10.1<sup>*</sup>; 10.4<sup>*</sup> | 0.6; 0.8 | 6.1; 1.9 | 0.5; 0.8 | 8.5<sup>*</sup>; 1.4 |
| main roof material | thatch / palm leaf | 0.5; 0.5 | 8.0<sup>*</sup>; 14.4<sup>*</sup> | 0.8; 0.7 | 0.7; 3.8<sup>*</sup> | 0.9; 0.7 | 0.2; 2.8 |
| tuberculosis can be cured | yes | 2.4; 2.2 | 6.9<sup>*</sup>; 11.8<sup>*</sup> | 1.8; 1.6 | 3.3; 4.9<sup>*</sup> | -; - | -; - |
| sons who have died | 1 | 1.7; 1.8 | 6.7<sup>*</sup>; 11.0<sup>*</sup> | 1.5; 1.3 | 3.8<sup>*</sup>; 2.0 | 1.6; 1.3 | 3.9; 1.2 |
| respondent's occupation (grouped) | sales | 1.7; 2.1 | 5.1<sup>*</sup>; 21.6<sup>*</sup> | 1.2; 1.3 | 0.7; 3.4<sup>*</sup> | 1.0; 1.3 | 0.0; 2.9 |
| had genital discharge in last 12 months | yes | 2.6; 2.6 | 4.1<sup>*</sup>; 8.6<sup>*</sup> | 2.1; 2.2 | 2.5; 6.1<sup>*</sup> | 2.0; 2.0 | 1.6; 3.3 |
| respondent's year of birth |  | 0.7; 0.6 | 15.2<sup>*</sup>; 51.6<sup>*</sup> | -; - | -; - | -; - | -; - |
| index last child prior to maternity-health (calendar) | no prior child | 0.4; 0.4 | 13.3<sup>*</sup>; 35.2<sup>*</sup> | 0.5; 0.8 | 5.6; 1.2 | 0.6; 1.0 | 2.9; 0.0 |
| total children ever born | 0 | 0.4; 0.4 | 13.1<sup>*</sup>; 22.3<sup>*</sup> | 0.8; 1.0 | 1.1; 0.0 | 0.9; 1.2 | 0.1; 0.5 |
| age at first sex | not had sex | 0.2; 0.2 | 12.2<sup>*</sup>; 21.3<sup>*</sup> | -; - | -; - | -; - | -; - |
| type of place of residence | rural | 0.5; 0.4 | 12.0<sup>*</sup>; 25.3<sup>*</sup> | -; - | -; - | -; - | -; - |
| fertility preference | no more | 1.8; 1.9 | 10.4<sup>*</sup>; 25.1<sup>*</sup> | 1.3; 1.1 | 1.7; 1.1 | 1.2; 1.1 | 1.0; 0.4 |
| sibling's date of birth (cmc) |  | 0.7; 0.7 | 10.2<sup>*</sup>; 28.3<sup>*</sup> | 1.0; 1.0 | 0.4; 0.1 | -; - | -; - |
| living children + current pregnancy | 0 | 0.5; 0.5 | 10.2<sup>*</sup>; 16.5<sup>*</sup> | 1.0; 1.4 | 0.1; 2.3 | 1.3; 1.6 | 0.8; 3.1 |
| ever been tested for hiv | yes | 1.8; 2.9 | 9.8<sup>*</sup>; 26.2<sup>*</sup> | -; - | -; - | -; - | -; - |
| wealth index factor score (5 decimals) |  | 1.4; 1.2 | 9.7<sup>*</sup>; 5.5<sup>*</sup> | 1.5; 1.0 | 1.5; 0.0 | 1.3; 1.1 | 0.6; 0.4 |
| fertility preference | have another | 0.6; 0.6 | 8.8<sup>*</sup>; 20.2<sup>*</sup> | 0.8; 1.0 | 1.8; 0.1 | 0.7; 1.1 | 2.2; 0.6 |
| sibling's date of birth (cmc) |  | 0.8; 0.7 | 8.6<sup>*</sup>; 24.3<sup>*</sup> | 1.0; 1.0 | 0.0; 0.2 | -; 0.6 | -; 1.6 |
| knows method | yes | 1.8; 2.5 | 8.5<sup>*</sup>; 11.2<sup>*</sup> | 1.2; 1.1 | 1.1; 0.5 | 1.2; 1.2 | 0.7; 0.4 |
| source for condoms: shop | yes | 1.7; 1.4 | 7.7<sup>*</sup>; 7.7<sup>*</sup> | 1.4; 1.2 | 3.2; 2.4 | 1.3; 1.2 | 1.4; 1.1 |
| knows method | yes | 1.7; 1.5 | 7.6<sup>*</sup>; 9.0<sup>*</sup> | 1.1; 0.8 | 0.8; 1.6 | 1.0; 0.8 | 0.1; 1.7 |
| source for condoms: pharmacy | yes | 1.9; 1.5 | 7.4<sup>*</sup>; 3.3<sup>*</sup> | 1.4; 1.1 | 2.5; 0.5 | 1.2; 1.1 | 0.7; 0.5 |
| source of drinking water | pipied to yard/plot | 2.1; 1.7 | 7.3<sup>*</sup>; 6.1<sup>*</sup> | 1.6; 1.4 | 3.0; 2.5 | 1.4; 1.5 | 1.3; 2.4 |
| wealth index | poorest | 0.4; 0.5 | 7.0<sup>*</sup>; 14.7<sup>*</sup> | -; - | -; - | -; - | -; - |
| knows method | yes | 1.8; 1.8 | 6.8<sup>*</sup>; 16.5<sup>*</sup> | 1.1; 1.1 | 0.5; 0.9 | 1.1; 1.0 | 0.3; 0.2 |
| sons elsewhere | 0 | 0.6; 0.5 | 6.7<sup>*</sup>; 19.2<sup>*</sup> | 0.8; 0.9 | 1.4; 1.2 | 0.9; 1.0 | 0.6; 0.2 |
| flag for v531 | no flag | 0.6; 0.7 | 6.4<sup>*</sup>; 5.5<sup>*</sup> | 0.8; 0.9 | 2.1; 1.0 | 0.7; 0.9 | 1.4; 0.5 |
| primary caregiver of children under age 18 | yes | 1.6; 1.6 | 6.4<sup>*</sup>; 7.2<sup>*</sup> | 1.1; 1.0 | 0.3; 0.2 | 1.1; 0.9 | 0.6; 0.5 |
| respondent's occupation | personal and protective services workers | 5.5; 2.7 | 6.1<sup>*</sup>; 8.3<sup>*</sup> | 3.6; 1.9 | 3.6; 3.5 | 4.0; 2.1 | 2.0; 3.6 |
| main floor material | earth, sand | 0.6; 0.6 | 6.0<sup>*</sup>; 8.4<sup>*</sup> | 0.9; 0.8 | 0.3; 1.6 | 1.0; 0.7 | 0.1; 1.9 |
| knows method | yes | 1.6; 2.0 | 6.0<sup>*</sup>; 6.3<sup>*</sup> | 1.2; 1.2 | 1.1; 0.9 | 1.0; 1.2 | 0.1; 0.4 |
| getting medical help for self: distance to health facility | not a big problem | 1.6; 1.4 | 6.0<sup>*</sup>; 6.3<sup>*</sup> | 1.2; 1.1 | 1.1; 0.6 | 1.1; 1.1 | 0.2; 0.6 |
| time to get to water source |  | 1.2; 1.2 | 5.9<sup>*</sup>; 5.1<sup>*</sup> | 1.1; 1.1 | 2.0; 1.5 | 1.1; 1.1 | 0.6; 1.5 |
| current marital status | never in union | 0.5; 0.5 | 5.7<sup>*</sup>; 17.3<sup>*</sup> | 0.9; 1.1 | 0.2; 0.6 | 0.9; 1.3 | 0.2; 1.2 |
| index last child prior to maternity-health (calendar) | 2 | 1.6; 1.4 | 5.7<sup>*</sup>; 5.8<sup>*</sup> | 1.3; 0.9 | 1.6; 0.4 | 1.1; 0.8 | 0.6; 2.0 |
| knows method | yes | 1.5; 1.4 | 5.5<sup>*</sup>; 7.3<sup>*</sup> | 1.0; 1.0 | 0.2; 0.0 | 1.0; 0.8 | 0.2; 1.5 |
| educational attainment | incomplete primary | 0.6; 0.7 | 5.3<sup>*</sup>; 5.2<sup>*</sup> | 0.8; 0.8 | 1.8; 3.6 | 0.9; 0.7 | 0.3; 2.2 |
| source of drinking water | public tap/standpipe | 1.5; 1.5 | 5.2<sup>*</sup>; 5.1<sup>*</sup> | 1.0; 1.1 | 0.1; 0.5 | 1.0; 1.0 | 0.1; 0.1 |
| tuberculosis spread by: don't know | yes | 0.6; 0.7 | 5.1<sup>*</sup>; 5.0<sup>*</sup> | 0.8; 0.9 | 1.4; 0.9 | 0.9; 1.0 | 0.3; 0.1 |
| knows method | yes | 1.6; 1.7 | 5.0<sup>*</sup>; 10.8<sup>*</sup> | 1.1; 1.1 | 0.7; 1.0 | 1.2; 1.1 | 0.7; 0.7 |
| age in 5-year groups | 30-34 | 1.7; 1.4 | 4.8<sup>*</sup>; 4.3<sup>*</sup> | 1.4; 1.1 | 2.1; 0.5 | 1.4; 1.1 | 1.5; 0.7 |
| sons elsewhere | 1 | 1.8; 1.9 | 4.7<sup>*</sup>; 10.7<sup>*</sup> | 1.4; 1.3 | 1.6; 1.7 | 1.1; 1.2 | 0.4; 0.7 |
| would want hiv infection in family to remain secret | yes | 1.4; 1.3 | 4.7<sup>*</sup>; 3.5<sup>*</sup> | 1.3; 1.2 | 2.9; 2.5 | 1.3; 1.1 | 2.4; 0.7 |
| cohabitation duration (grouped) | 15-19 | 1.6; 1.7 | 4.6<sup>*</sup>; 8.8<sup>*</sup> | 1.3; 1.2 | 1.6; 1.5 | 1.2; 1.1 | 0.7; 0.7 |
| don't know any source for condoms | yes: no source known | 0.6; 0.4 | 4.3<sup>*</sup>; 8.6<sup>*</sup> | 0.8; 0.7 | 0.6; 1.5 | 1.1; - | 0.2; - |
| heard family planning on tv last few months | yes | 1.5; 1.6 | 3.8<sup>*</sup>; 8.9<sup>*</sup> | 1.0; 1.2 | 0.0; 2.1 | 0.9; 1.3 | 0.2; 2.3 |
| literacy | cannot read at all | 0.7; 0.8 | 3.8<sup>*</sup>; 4.1<sup>*</sup> | 0.8; 0.8 | 0.8; 2.2 | 0.9; 0.9 | 0.2; 0.7 |
| getting medical help for self: not wanting to go alone | not a big problem | 1.6; 1.6 | 3.4<sup>*</sup>; 5.6<sup>*</sup> | 1.2; 1.2 | 0.9; 1.4 | 1.1; 1.1 | 0.4; 0.4 |
| willing to care for relative with aids | yes | 2.8; 2.3 | 3.4<sup>*</sup>; 4.7<sup>*</sup> | 2.0; 1.6 | 1.8; 1.5 | 1.8; 2.2 | 0.8; 2.0 |
| drugs to avoid hiv transmission to baby during pregnancy | yes | 1.6; 1.9 | 3.4<sup>*</sup>; 5.9<sup>*</sup> | 1.2; 1.4 | 0.7; 1.8 | -; - | -; - |
| source of drinking water | unprotected well | 0.5; 0.7 | 3.4<sup>*</sup>; 3.4<sup>*</sup> | 0.8; 0.9 | 0.5; 0.9 | 1.1; 1.0 | 0.1; 0.0 |
| living children + current pregnancy | 3 | 1.5; 1.6 | 3.4<sup>*</sup>; 7.7<sup>*</sup> | 1.3; 1.3 | 1.7; 3.0 | 1.2; 1.1 | 0.9; 0.4 |
| knows method | yes | 1.9; 2.3 | 3.4<sup>*</sup>; 4.3<sup>*</sup> | 0.9; 0.8 | 0.1; 0.5 | 0.8; 0.6 | 0.5; 1.0 |
| respondent's occupation | sales and services elementary occupations | 1.6; 2.1 | 3.4<sup>*</sup>; 6.8<sup>*</sup> | 1.1; 1.6 | 0.2; 2.6 | 0.9; 1.4 | 0.2; 1.4 |
| tuberculosis spread by: air when coughing or sneezing | yes | 1.4; 1.4 | 3.1<sup>*</sup>; 5.0<sup>*</sup> | 1.1; 1.2 | 0.4; 1.3 | 1.0; 1.0 | 0.1; 0.0 |
| main wall material | cement | 1.6; 1.4 | 3.0<sup>*</sup>; 3.9<sup>*</sup> | 1.4; 1.3 | 1.3; 1.7 | 1.2; 1.2 | 0.7; 1.3 |
| living children + current pregnancy | 1 | 1.7; 1.3 | 7.8<sup>*</sup>; 2.0 | 1.6; 1.5 | 5.3<sup>*</sup>; 4.7<sup>*</sup> | 1.4; 1.5 | 2.4; 3.9<sup>*</sup> |
| living children + current pregnancy (grouped) | 6+ | 0.5; 0.6 | 4.7; 9.6<sup>*</sup> | 0.3; 0.3 | 13.3<sup>*</sup>; 34.2<sup>*</sup> | 0.3; 0.2 | 6.2; 23.4<sup>*</sup> |
| births in last three years | 0 | 1.4; 1.7 | 3.6; 19.1<sup>*</sup> | 1.6; 1.9 | 5.6<sup>*</sup>; 22.6<sup>*</sup> | 1.5; 2.0 | 3.2; 14.7<sup>*</sup> |
| births in last three years | 1 | 0.8; 0.6 | 2.6; 14.0<sup>*</sup> | 0.7; 0.6 | 4.5<sup>*</sup>; 17.2<sup>*</sup> | 0.7; 0.5 | 2.6; 13.0<sup>*</sup> |
| births in last five years | no births | 1.2; 1.4 | 1.3; 8.6<sup>*</sup> | 1.5; 1.9 | 3.8<sup>*</sup>; 19.5<sup>*</sup> | 1.5; 2.0 | 2.6; 14.7<sup>*</sup> |
| relationship to household head** | wife | 0.8; 0.8 | 1.0; 3.3<sup>*</sup> | 0.6; 0.5 | 5.7<sup>*</sup>; 23.1<sup>*</sup> | 1.0; 0.5 | 0.0; 18.4<sup>*</sup> |
| ideal number of children | 7 | 0.3; 0.5 | 4.0<sup>*</sup>; 2.8 | 0.3; 0.5 | 3.1<sup>*</sup>; 4.3<sup>*</sup> | 0.3; 0.4 | 2.1; 3.3 |
| rohrer's index |  | 0.9; 0.9 | 0.6; 0.9 | 0.7; 0.7 | 6.0<sup>*</sup>; 9.9<sup>*</sup> | 0.8; 0.7 | 4.6<sup>*</sup>; 9.4<sup>*</sup> |
| daughters at home | 0 | 1.1; 1.0 | 0.4; 0.1 | 1.6; 1.9 | 4.3<sup>*</sup>; 17.8<sup>*</sup> | 1.8; 1.9 | 4.8<sup>*</sup>; 15.0<sup>*</sup> |
| body mass index |  | 1.0; 1.0 | 0.1; 0.1 | 0.8; 0.8 | 6.2<sup>*</sup>; 10.1<sup>*</sup> | 0.8; 0.7 | 4.7<sup>*</sup>; 9.5<sup>*</sup> |
| current marital status | married | 0.8; 0.8 | 1.7; 2.6 | 0.5; 0.5 | 7.3<sup>*</sup>; 24.6<sup>*</sup> | 0.9; 0.5 | 0.1; 19.6<sup>*</sup> |
| currently/formerly/never in union | currently in union/living with a man | 0.8; 0.8 | 1.1; 2.3 | 0.5; 0.5 | 6.0<sup>*</sup>; 24.4<sup>*</sup> | 1.1; 0.5 | 0.2; 20.0<sup>*</sup> |
| sons at home | 0 | 1.1; 1.0 | 0.3; 0.2 | 1.6; 1.8 | 5.3<sup>*</sup>; 15.8<sup>*</sup> | 1.7; 2.0 | 5.1; 15.4<sup>*</sup> |
| daughters at home | 3 | 0.7; 0.8 | 1.5; 1.3 | 0.5; 0.6 | 3.2<sup>*</sup>; 5.8<sup>*</sup> | 0.5; 0.5 | 2.5; 4.3 |
| age in 5-year groups | 45-49 | 0.9; 1.2 | 0.4; 1.1 | 0.4; 0.4 | 6.0<sup>*</sup>; 9.0<sup>*</sup> | 0.4; 0.5 | 2.7; 4.6 |
| number of children 5 and under in household (de jure) | 7 | 1.1; 0.6 | 0.0; 0.2 | 2.1; 0.6 | 0.2; 0.2 | 0.0; 0.0 | 22.8<sup>*</sup>; 55.5<sup>*</sup> |

\* Replicated in analysis. Specifically, these variables were above the false discovery rate 5% threshold in the discovery dataset, and had a nominal p-value <0.05 in the replication dataset.
\*\* Variables indicated with double asterisk were used as controls in the *ex ante* and/or *post* analysis, and were tested in a model with only the *ex post* variables as independent predictors.

Several variables stand out for their association with HIV-1 and for raising potentially useful targets for future investigation. Three variables were associated with HIV-1 in all three analyses *and* both surveys: having exactly 1 birth in the past 5 years (increased risk), currently breastfeeding (decreased risk), and desiring to delay having children for more than two years (decreased risk). Eleven additional variables were associated with HIV-1 positivity in all but one of the *ex post* analyses (including several variables that were used as *ex post* control): being formerly and not currently married, including divorce and widowhood (3 variables, all increase risk), variables related to being the head of the household (3 variables, all increase risk), the number of children (3 variables, fewer children confer increased risk), currently using a condom for contraception (increased risk), and an indicator for ownership of a bicycle in the household (decreased risk).

Figure 3 shows the sub-group associations for the variables that were replicated in univariate analyses in the overall sample in both surveys and in at least 17 out of the 18 subgroups examined (9 subgroups, as noted above, in each of the 2 surveys). A total of 8 variables met these criteria (including several that were associated with HIV-1 in all full-sample analyses, denoted with *): the indicators for widowhood and being formerly in a union*, being the head of the household*, having exactly 2 people in the household (relative to all other household sizes), age, reporting a genital ulcer in the past 12 months, owning a bicycle*, and currently breastfeeding*.

**Figure 3.**
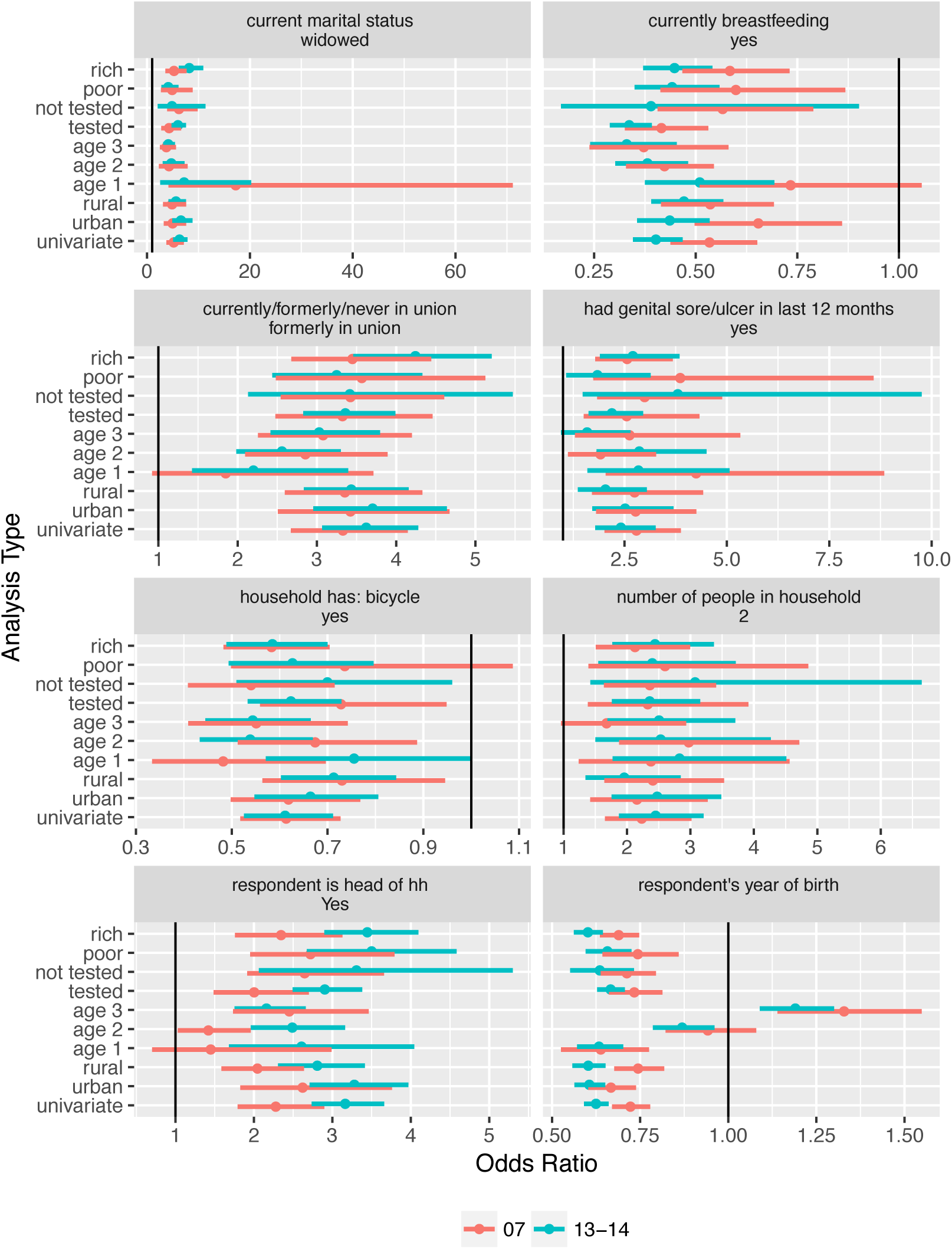
Strength of association for univariate overall and population subgroups. Stratas include wealth index less than or equal to 3 and greater than 3 (“poor” and “rich” respectively), individuals that have had or have not had an HIV-1 test (“tested” and “not tested”), individuals living a rural (“rural”) or urban (“urban”) areas, and of ages less than 23 (“age1”), between 23 and 33 (“age2”), and older than 33 (“age3”).

Two general categories of variables were associated with HIV-1 in the *ex ante* and *ex post* analyses but not in the univariate analyses (i.e. variables whose association with HIV-1 was “uncovered” after adjustment, shown at the bottom variables in Table 2). These include variables related to the number of children that were different from those replicated in all or nearly all analyses (again, fewer children confer increased risk), and the anthropometric measurements body-mass index and Rohrer’s index (higher index associated with lower HIV-1 risk).

Figure 4 shows the extent to which the variables that were replicated in the *ex ante* analysis are correlated and clustered among themselves. We observed that the correlation pattern between the 2007 and 2013-2014 surveys were strikingly similar. We hypothesized that this may be partly a reflection of the variable construction process (e.g. being married would be expected to be strongly negatively correlated with being divorced), and partly of the likely stable social and environmental patterns in Zambia over this time period (Supplementary Figures SA1-SA2).

**Figure 4.**
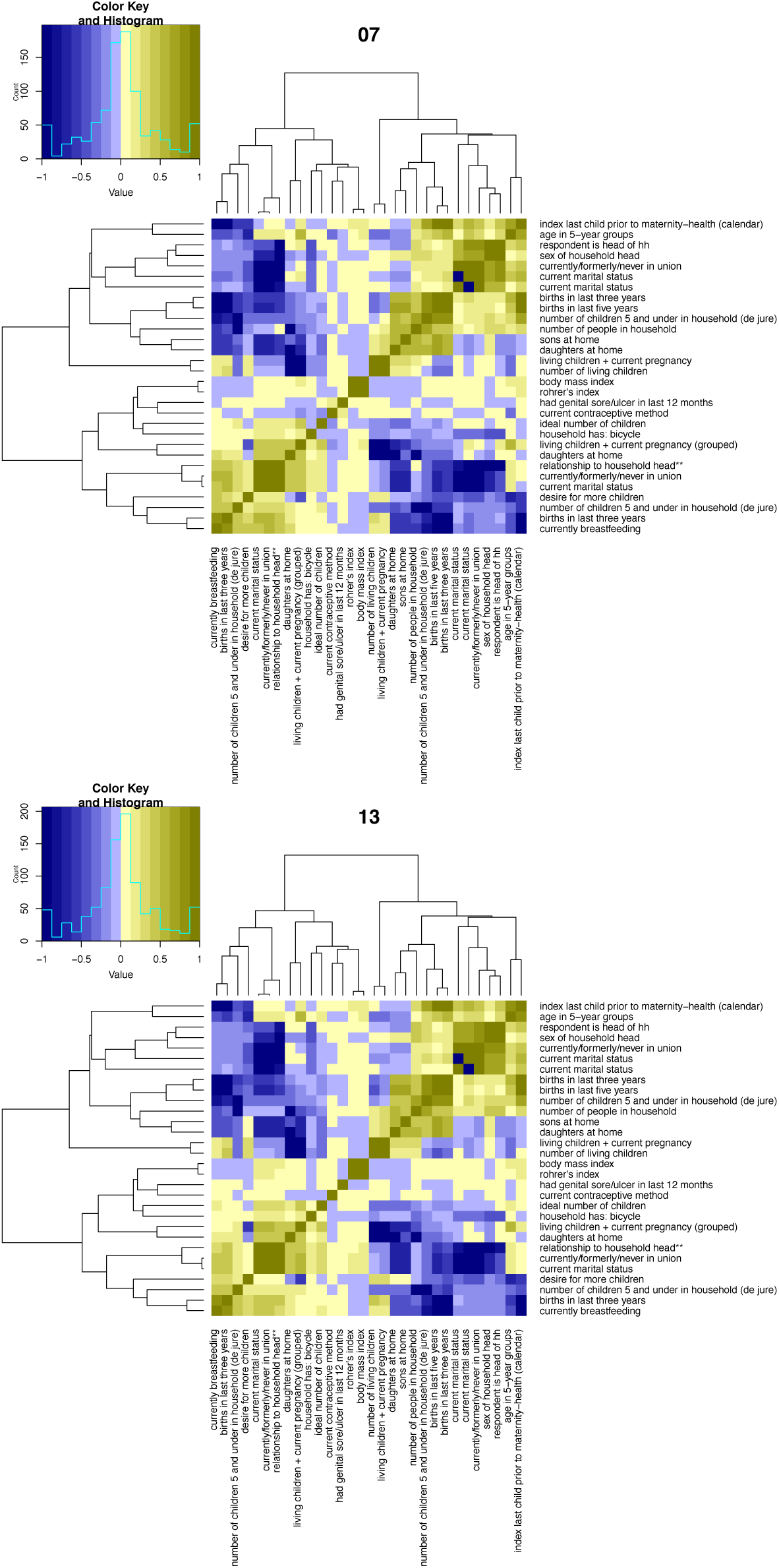
Correlation matrix of variables replicated in the *ex ante* analysis. Top panel shows results for the 2007 survey and the bottom panel for the 2013-2014 survey. Variable clusters representing similar constructs appear in both surveys, such as variables that characterize marital status.

In the three analyses and the two surveys, the variance explained in HIV-1 status ranged from 0.21 (in the *ex post* analysis of the 2007 survey) to 0.32 (in the univariate analysis of the 2013-2014 survey). The area under the curve in the six analyses ranged from 0.76 to 0.82 (specificity=0.75 at sensitivity=0.75) (Supplementary Figure SA3).

## Discussion

We describe the findings from the first XWAS of HIV-1 risk in nationally representative sero-surveys. Out of all the variables tested (688 in 2007 and 727 in 2013-2014, of which 688 were overlapping), we identify several candidate variables that are associated with HIV-1 when examined in multiple analyses and may present opportunities for identifying previously under-recognized risks. These include positive associations with widowhood/divorce/being formerly in union, being the head of the household, having a small household size, and reporting a genital ulcer in the past 12 months; and negative associations with breastfeeding and bicycle ownership. The reasons for these consistent associations may have different implications. A causal relationship may have implications for targeting and design of prevention interventions. A non-causal relationship (that is observed due to confounding or reverse causation) may still have benefits for testing programs that are interested in increasing testing among high-prevalence groups. The nature of the associations, therefore, deserves further discussion.

The reason for the positive association of widowhood with HIV-1 may be due to widows’ engagement in high-risk behaviors for basic income and sustenance; it may also be partly caused by HIV-1 positivity among the widows’ now-deceased husbands. Our study cannot tease apart the dominant causal pathway, and both may contribute to the association. The similar effect among divorced women is more consistent with risky behaviors following the loss of a spouse. Recent evidence also supports a causal role: a nationally representative survey of HIV-1 incidence in Rwanda from 2013-2014 found elevated rate of new infections in widows.^29^ If widowhood and divorce lead to increased HIV-1 risk, then targeting of prevention interventions such as pre-exposure prophylaxis may mitigate the associated risk. If HIV-1 prevalence is higher among these women because of pre-existing risk, then this finding may still help in guiding HIV-1 identification for early treatment and care that may reduce their risk of infecting others.

The relationship between current breastfeeding and HIV-1 risk is also notable. It is not a known factor that decreases risk of HIV-1 acquisition.^30,31^ This association may indicate the decreased propensity to breastfeed among HIV-positive women. While public health guidelines for breastfeeding among HIV-infected women has shifted over the past decade, breastfeeding has been recommended by the World Health Organization since 2010.^32,33^ Since we find decreased risk of HIV-1 among women who breastfeed in both 2007 and 2013-2014 (in all three analyses and 17 subgroups), this finding may indicate the challenges of changing breastfeeding behaviors and the importance of finding effective approaches to behavior change in this domain.

The variables that we highlight were replicated in multiple analyses, but this study also identifies factors whose less consistent association with HIV-1 may nevertheless warrant additional consideration. Several variables related to method of contraception were positively associated with HIV-1 status in the univariate analyses, including hormonal contraceptives, condoms, and female condoms. Wealth was associated with HIV-1 in the univariate analysis (higher risk among wealthier women), but not in the adjusted models. No variable identifying educational attainment was associated with HIV-1 in the adjusted models. These assessments improve on the extant assessment of epidemiologic risk that are commonly presented along with the DHS data (and commonly used by UNAIDS and others).^4,13^ The DHS stratifies HIV-1 risk by age, residence, marital status, education, and wealth. XWAS improves on such stratifications by reducing potential bias from failure to consider other relevant covariates, and by using an FDR that accounts for multiple comparisons.

The extent to which our findings are generalizable to other contexts is unknown. Extending HIV-1 XWAS to additional surveys across sub-Saharan Africa and over time, however, is readily feasible and will enable greater understanding of the generalizability and stability over time of our findings. We note that the putative variables we identified in common in the 2007 and 2013-2014 surveys had similar association sizes in both surveys. These similar association sizes and correlations points to the stability of social, behavior, and environment over time in Zambia.

Some limitations deserve explicit mention. First, we only tested variables with at least 90% complete data. While we retained approximately 700 variables for analysis, some important variables could have been excluded because of missingness. Second, the error rate among self-reported variables may also bias results. Errors are more likely for some variables than for others. Any non-differential bias (e.g., individuals that report inaccurately in both HIV-1 positive and negative individuals) will lead to loss of power and correlations that are closer to null; however, we emphasize that sample sizes in our investigation are large. Third, self-reported variables may exhibit differential bias if participants answer differentially based on HIV-1 status. Differential bias may distort effects in unpredictable ways. Fourth, we could not assess association with incident or recent infection to mitigate chances of reverse causality. While some DHS surveys also measure CD4 cell counts (that may proxy for duration of infection), the Zambia surveys did not, and we did not control for duration of infection (except through some indirect controlling by adjusting for age). It is plausible, for example, that a decrease in body-mass index is a consequence of HIV-1 rather than a cause. Challenges to causal identification are a generic issue in large-scale cross-sectional association studies, but such analyses nevertheless remain an important method to identify potential risk factors.^34^

In summary, we report the findings from the first XWAS of HIV-1 risk from nationally representative surveys of social, economic, environmental, and behavioral factors in Zambia. We identify strong and consistent associations with widowhood, breastfeeding, and several other self-reported indicators that may be amenable to further investigations and interventions and that may be used to guide screening policies.

## Author Contributions

EB and CJP conceived the work and carried out the analyses. JB and JPAI critically assessed the methods and findings, and contributed to the study conceptualization and preparation of the manuscript.

## Role of the Funding Organization or Sponsor

The sponsor had no role in the design, interpretation or conclusions of this study.

## Previous Presentations

An early version of this work was presented at the Conference on Retroviruses and Opportunistic Infections in Boston, 2016.

## References

1. 90–90–90 - An ambitious treatment target to help end the AIDS epidemic. (Accessed December 10, 2015, at http://www.unaids.org/en/resources/documents/2014/90-90-90.)

2. Deeks SG, Lewin SR, Havlir DV. The end of AIDS: HIV infection as a chronic disease. The Lancet 2013; 382: 1525–33.

3. Staveteig S, Bradley S, Nybro E, Wang S. Demographic Patterns of HIV Testing Uptake in Sub-Saharan Africa. DHS Comparative Reports No. 30. ICF International.2013.

4. UNAIDS AIDSinfo: Epidemiological Status. (Accessed April 31, 2016, at http://aidsinfo.unaids.org/.)

5. UNAIDS 2014 Gap Report. (Accessed August 31, 2015, at http://www.unaids.org/en/resources/documents/2014/20140716_UNAIDS_gap_report.)

6. Demographic and Health Surveys. ICF International. (Accessed August 3, 2015, at http://www.measuredhs.com.)

7. Opening Statement From Ambassador Deborah L. Birx, M.D., at the UNAIDS 37th Programme Coordinating Board Meeting. (Accessed August 10, 2016, at http://www.pepfar.gov/press/releases/2015/248739.htm.)

8. PEPFAR’s Dr Deborah Birx urges sharper focus to halt HIV globally. (Accessed August 20, 2016, at https://www.fic.nih.gov/News/GlobalHealthMatters/january-february-2016/Pages/deborah-birx-pepfar-global-hiv-control.aspx).

9. WHO HIV/AIDS Strategic Information: Surveillance. (Accessed August 31, 2015, at http://www.who.int/hiv/strategic/surveillance/en/.)

10. De Cock KM, Rutherford GW, Akhwale W. Kenya AIDS Indicator Survey 2012. JAIDS Journal of Acquired Immune Deficiency Syndromes 2014;66:S1–S2.

11. Anderson S-J, Cherutich P, Kilonzo N, et al. Maximising the effect of combination HIV prevention through prioritisation of the people and places in greatest need: a modelling study. The Lancet 2014; 384: 249–56.

12. Zambia DHS, 2007 - Final Report. (Accessed June 13, 2016, at http://dhsprogram.com/publications/publication-FR211-DHS-Final-Reports.cfm.)

13. Zambia DHS, 2013-14 - Final Report. (Accessed August 31, 2015, at http://dhsprogram.com/publications/publication-FR304-DHS-Final-Reports.cfm.)

14. Patel CJ, Ioannidis JP. Studying the elusive environment in large scale. J Am Med Assoc 2014; 311: 2173–4.

15. Ioannidis J. Why Most Published Research Findings Are False. PLoS Med 2005; 2: e124.

16. Tzoulaki I, Patel CJ, Okamura T, et al. A nutrient-wide association study on blood pressure. Circulation 2012; 126: 2456–64.

17. Patel CJ, Rehkopf DH, Leppert JT, et al. Systematic evaluation of environmental and behavioural factors associated with all-cause mortality in the United States National Health and Nutrition Examination Survey. Int J Epidemiol 2013; 42: 1795–810.

18. Patel CJ, Cullen MR, Ioannidis JP, Butte AJ. Systematic evaluation of environmental factors: persistent pollutants and nutrients correlated with serum lipid levels. Int J Epidemiol 2012; 41: 828–43.

19. Patel CJ, Bhattacharya J, Butte AJ. An environment-wide association study (EWAS) on type 2 diabetes mellitus. PloS one 2010; 5: e10746.

20. Ioannidis JP, Tarone R, McLaughlin JK. The false-positive to false-negative ratio in epidemiologic studies. Epidemiology 2011; 22: 450–6.

21. Patel CJ, Ioannidis JP. Placing epidemiological results in the context of multiplicity and typical correlations of exposures. Journal of epidemiology and community health 2014;68:1096–100.

22. Dong Y, Peng C-YJ. Principled missing data methods for researchers. SpringerPlus 2013; 2: 1.

23. Benjamini Y, Hochberg Y. Controlling the false discovery rate: a practical and powerful approach to multiple testing. Journal of the royal statistical society Series B (Methodological) 1995: 289–300.

24. White H. A heteroskedasticity-consistent covariance matrix estimator and a direct test for heteroskedasticity. Econometrica: Journal of the Econometric Society 1980: 817–38.

25. Rutstein S, Johnson K. The DHS Wealth Index. DHS Comparative Reports No. 6. Calverton, Maryland: ORC Macro; 2004. 2011.

26. Yang J, Ferreira T, Morris AP, et al. Conditional and joint multiple-SNP analysis of GWAS summary statistics identifies additional variants influencing complex traits. Nature genetics 2012; 44: 369–75.

27. Patel CJ, Cullen MR, Ioannidis JPA, Rehkopf DH. Systematic assessment of the correlation of household income with infectious, biochemical, physiological factors in the United States, 1999-2006¨ Am J Epidemiol 2014; 181: 171–9.

28. Johnson RA, Wichern DW. Applied multivariate statistical analysis: Prentice hall Englewood Cliffs, NJ; 1992.

29. Remera E, Kanters S, Mulidabigwi A, et al. 2013-14 Rwanda HIV Incidence Household Survey: Understanding HIV Epidemic in Rwanda. CROI 2016. Boston, USA.

30. Serwadda D, Wawer MJ, Musgrave SD, Sewankambo NK, Kaplan JE, Gray RH. HIV risk factors in three geographic strata of rural Rakai District, Uganda. Aids 1992; 6: 983–90.

31. Cain D, Simbayi L, Kalichman S, Cherry C, Jooste S, Mfecane S. Risk factors for HIV-AIDS among youth in Cape Town, South Africa. 2015.

32. World Health Organization. HIV and Infant Feeding: Update. 2006. (Accessed November 17, 2016, at http://apps.who.int/iris/bitstream/10665/43747/1/9789241595964_eng.pdf.)

33. World Health Organization. Guidelines on HIV and Infant Feeding: Principles and recommendations for infant feeding in the context of HIV and a summary of evidence. 2016. (Accessed November 17, 2016, at https://www.unicef.org/aids/files/hiv_WHO_guideline_on_HIV_and_IF.pdf.)

34. Ioannidis J. Exposure-wide epidemiology: revisiting Bradford Hill. Statistics in medicine 2015.

